# Structural characterization of HIV-1 Env heterotrimers bound to one or two CD4 receptors reveals intermediate Env conformations

**DOI:** 10.1101/2023.01.27.525985

**Authors:** Kim-Marie A. Dam, Chengcheng Fan, Zhi Yang, Pamela J. Bjorkman

**Affiliations:** Division of Biology and Biological Engineering, California Institute of Technology, Pasadena, CA, USA; Department of Molecular and Cell Biology, University of California, 13 Berkeley, CA 94720, USA

## Abstract

HIV-1 envelope (Env) exhibits distinct conformational changes in response to host receptor (CD4) engagement. Env, a trimer of gp120/gp41 heterodimers, has been structurally characterized in a closed, prefusion conformation with closely associated gp120s and coreceptor binding sites on gp120 V3 hidden by V1V2 loops, and in fully-saturated CD4-bound open Env conformations with changes including outwardly rotated gp120s and displaced V1V2 loops. To investigate changes resulting from sub-stoichiometric CD4 binding, we solved 3.4Å and 3.9Å single-particle cryo-EM structures of soluble, native-like Envs bound to one or two CD4 molecules. Env trimer bound to one CD4 adopted the closed, prefusion Env state. When bound to two CD4s, the CD4-bound gp120s exhibited an open Env conformation including a four-stranded gp120 bridging sheet and displaced gp120 V1V2 loops that expose the coreceptor sites on V3. The third gp120 adopted an intermediate, occluded-open state that included gp120 outward rotation but maintained the prefusion, three-stranded gp120 bridging sheet and showed only partial V1V2 displacement and V3 exposure. We conclude that engagement of one CD4 molecule was insufficient to stimulate CD4-induced conformational changes, while binding two CD4 molecules led to Env opening in CD4-bound protomers only. Together, these results illuminate HIV-1 Env intermediate conformations and illustrate the structural plasticity of HIV-1 Env.

---

The HIV-1 envelope (Env) glycoprotein, a heavily glycosylated homotrimer containing gp120 and gp41 subunits, mediates entry into host cells to initiate infection^1^. On the surface of virions, Env adopts a closed, prefusion conformation similar to that observed in soluble, native-like Env trimer ectodomains^2–5^. The viral entry process is initiated when gp120s bind to the host receptor, CD4, at the CD4-binding site (CD4bs) located distal to the Env apex on the sides of each of the three gp120s^6–10^. This triggers conformational changes in gp120 that expose the gp120 V3 coreceptor binding site that is occluded in the prefusion conformation beneath gp120 V1V2 loops^6–10^. Coreceptor binding results in further conformational changes that lead to insertion of the gp41 fusion peptide into the host cell membrane and fusion of viral and host membranes^1,10^.

X-ray crystallography and single particle cryo-electron microscopy (cryo-EM) structures have characterized soluble versions of HIV-1 Envs^11^ in closed, prefusion^2,3^, CD4-bound open^6,7,10^, and intermediate, partially-open conformations^6,8,12^. Multiple studies have demonstrated that the native-like soluble Envs (SOSIPs)^11^ used for structural studies resemble virion-bound Envs, suggesting these conformations are relevant to the viral Env entry process^4,5,11,13–15^. The closed, prefusion Env conformation is characterized by gp120 V1V2 loops interacting around the trimer apex, thereby shielding the coreceptor binding sites on the V3 loops^2,3,16^. Structures of CD4-bound open Env trimers revealed receptor-induced changes in which the gp120 subunits rotated outwards, the V1V2 loops were displaced from the apex by ~40 Å to the sides of Env, and the coreceptor binding site on each V3 was exposed and became mostly disordered^6–10^ (Supplementary Movie 1). This process also converted the closed, prefusion conformation three-stranded gp120 bridging sheet composed of the β20, β21, and β3 β-strands^2^ to a four-stranded antiparallel β-sheet in which strand β2, whose residues are located in a proximal helix in the closed prefusion formation, intercalated between strands β21 and β3^2,6–8^. Intermediate Env conformations include occluded open^6,12^ and partially open conformations^8,17^. In the occluded open conformation observed in trimer complexes with the CD4bs antibody b12^6^ and with similar antibodies raised in vaccinated non-human primates^12^, the gp120 subunits were outwardly rotated from the central trimer axis as in CD4-bound open conformations, but V1V2 displacement and V3 exposure did not occur and the prefusion three-stranded gp120 sheet was maintained^6,12^. In partially open Env conformations, CD4 binding led to the characteristic CD4-induced structural changes in gp120 but subsequent binding of the gp120-gp41 interface antibody 8ANC195 led to partial closure of the gp120s^8^.

A prevailing enigma regarding Env conformational changes and the role of CD4 in initiating the fusion process concerns whether the gp120/gp41 protomers that form the Env trimer behave cooperatively or independently during receptor-induced transformations. This information would reveal how many CD4 receptor and CCR5 coreceptor molecules are needed to engage each Env trimer to induce fusion and further elucidate Env function as it relates to virus infectivity, thereby informing the design of entry inhibitors and mechanisms of antibody neutralization and fusion. To illuminate the role of receptor stoichiometry in CD4-induced conformational changes in HIV-1 Env, we designed soluble Env heterotrimers that can bind only one or only two CD4 receptors for comparisons with Env homotrimers binding either zero CD4s (closed, prefusion trimers) or three CD4s (fully-saturated CD4-bound open trimers). Using single-particle cryo-EM, we solved structures of one or two CD4s bound to the clade A BG505 trimer^11^ to 3.4 and 3.9 Å, respectively. We found that binding one CD4 resulted in a closed, prefusion Env conformation that showed only subtle indications of CD4-induced changes. Binding two CD4 molecules induced an asymmetric, partially open Env conformation in which the gp120 subunits resembled open (for the CD4-bound protomers) and occluded-open (for the unliganded protomer) conformations, while the three gp41 subunits were structurally different from each other. Together, these results illustrate intermediate Env conformations and inform our understanding of the events that lead to HIV-1 fusion.

## Results

### Design and validation of heterotrimer Env constructs

A soluble heterotrimer Env that can bind only one CD4 receptor, termed HT1, was generated by co-expressing plasmids encoding BG505 SOSIP.664^11,18^ bearing a D368R mutation that eliminates CD4 binding^19,20^ and an affinity-tagged mutant BG505 SOSIP.664 at a 20:1 ratio (Extended Data Fig. 1a). For HT2, which binds only two CD4 receptors, plasmids encoding BG505 SOSIP.664 and a tagged BG505-D368R SOSIP.664 were co-expressed in a 20:1 ratio (Extended Data Fig. 1a). Assuming random assembly, 13% of the Env population would be composed of the desired singly-tagged heterotrimer and less than 1% would contain dually- and triply-tagged trimers^21^. For both constructs, immunoaffinity column purification resulted in purified tagged heterotrimers (Extended Data Fig. 1a).

To validate the design and purification of BG505-HT1 and BG505-HT2, we performed enzyme-linked immunosorbent assays (ELISAs) to compare binding of soluble CD4 to heterotrimeric Envs and to homotrimeric BG505 (both wildtype and D368R mutant) Envs (Extended Data Fig. 1b). As expected, wildtype BG505 exhibited the highest level of CD4 binding, BG505-D368R showed only limited CD4 binding at high concentrations, and BG505-HT1 and -HT2 showed intermediate levels of CD4 binding, with more binding to -HT2 than to -HT1.

### BG505-HT1 bound to one CD4 adopts a closed, prefusion Env trimer conformation

We used single-particle cryo-EM to solve a 3.4 Å structure of BG505-HT1 heterotrimer bound to one CD4 molecule (Fig. 1a, Extended Data Fig. 2). Despite CD4 recognition of the CD4bs of one gp120 protomer, the Env trimer maintained the prefusion, closed conformation with V1V2 loops at the Env apex and V3 loops shielded beneath V1V2^2,3,11^ (Fig. 1b; Supplementary Movie 1), suggesting that interactions of a soluble Env trimer with one CD4 molecule are insufficient to trigger conformational changes that lead to Env opening^6–8^. The ability of CD4 to bind to a closed Env conformation was previously observed in a low-resolution structure of CD4 bound to a homotrimeric SOSIP that included mutations to prevent Env opening^22^, whereas the HT1 heterotrimer used for the structural studies reported here did not include mutations that lock Env into a closed, prefusion conformation.

**Fig. 1:**
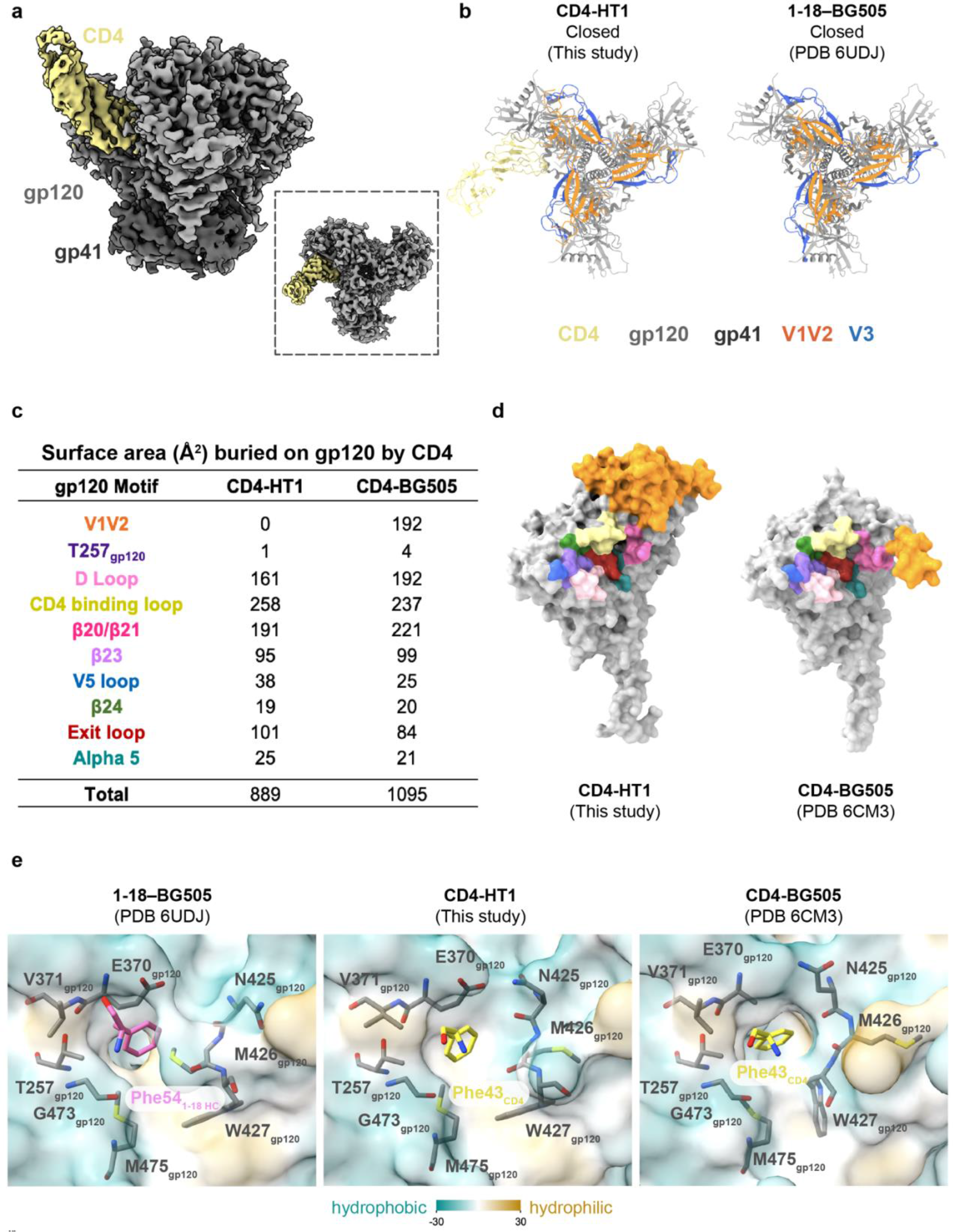
3.4 Å cryo-EM structure of BG505-HT1 bound to one CD4 reveals closed, pre-fusion Env conformation. **a**, Side view of the 3.4 Å CD4-HT1 density map. Inset: top-down view. **b**, Top-down cartoon representations of CD4-HT1 and 1-18–BG505 (PDB 6UDJ) structures with gp120 V1V2 and V3 loops highlighted. **c**, Table summarizing BSA on gp120 from CD4 binding for CD4-HT1 and CD4-BG505 (PDB 6CM3) complexes. **d**, Surface representation comparisons of CD4-HT1 and CD4-BG505 (PDB 6CM3). **e**, Surface representations depicting hydrophobicity (Kyte-Doolittle scale^57^) for 1-18–BG505 (PDB 6UDJ), CD4-HT1, and CD4-BG505 (PDB 6CM3) overlaid with stick representations of gp120 residues within the Phe43 cavity.

We next compared interactions in the CD4bs of the CD4-bound protomer of the CD4-HT1 complex with the CD4bs in the gp120 of a CD4-bound open BG505 trimer (PDB 6CM3) by calculating the surface area on gp120 buried by CD4 (buried surface area; BSA) (Fig. 1c,d). The BSA within the CD4bs was comparable for CD4-HT1 and CD4-BG505 gp120s but for gp120s in the CD4-BG505 Env complex, V1V2 displacement led to an additional ~200 A^2^ of BSA on gp120 (Fig. 1c,d), which was previously shown to stabilize the CD4-induced, open Env conformation^6,7^. These comparisons suggest that the Env-CD4 interface remains largely unchanged during CD4 engagement, the primary difference in the CD4-bound open structure is the displacement of V1V2 from the Env apex to the side of gp120 where it makes additional contacts with CD4.

CD4-induced Env conformational changes are triggered, at least in part, by insertion of Phe43_CD4_ into a conserved, hydrophobic cavity (the Phe43 cavity) on gp120^6–8,23,24^. Small molecule CD4 mimetics such as BNM-III-170 and M48U1 insert hydrophobic entities into the Phe43_CD4_ cavity, thereby competing with CD4 binding and inducing Env opening^9,25–30^. Some CD4bs broadly neutralizing antibodies (bNAbs) also mimic Phe43_CD4_ interactions by inserting a hydrophobic residue at antibody heavy chain (HC) position 54 into the Phe43 cavity on gp120. However, by contrast to the conformational effects of CD4 and selected small mimetic inhibitors on Env conformation, CD4bs bNAbs with a hydrophobic HC residue 54 stabilize the prefusion, closed Env conformation when bound to trimeric Env^31–35^.

To examine the consequences of insertion of Phe43_CD4_ into a single gp120 Phe43 cavity in the CD4-HT1 complex, we compared the structural landscape of the Phe43 cavity in the gp120s of two symmetric Env trimer complexes: the CD4bs bNAb 1-18 bound to a closed, prefusion conformation BG505^33^ and CD4 bound to an open, fully-saturated CD4-bound BG505 trimer^8^ (Fig. 1e). We identified and compared the positions of conserved residues in the region, some of which have been characterized to undergo rearrangements during CD4-induced Env opening^7,23^. Residues in the CD4 binding loop (E370_gp120_, V371_gp120_) along with T257_gp120_ and exit loop (G473_gp120_, M475_gp120_) residues maintained analogous positions in the one CD4-bound HT1, zero CD4-bound closed, and three CD4-bound open trimers (Fig. 1e). However, subtle differences in the gp120 β20/β21 loop were observed. For example, in the 1-18–BG505 complex, the N425_gp120_ side chain pointed away from Phe54_1-18 HC_, while the M426_gp120_ side chain pointed towards Phe54_1-18 HC_ and the planes of the W427_gp120_ side chain and Phe54_1-18 HC_ side chain were parallel. By contrast, in the CD4-BG505 open complex, the N425_gp120_ side chain pointed upward from the Phe43 cavity ceiling, the M426_gp120_ side chain pointed away from Phe43_CD4_, and the W427_gp120_ side chain was perpendicular to the Phe43_CD4_ side chain. The CD4-HT1 complex showed an intermediate orientation of gp120 β20/β21 loop residues, with the N425_gp120_ and M426_gp120_ side chains adopting positions similar to their positions in the CD4-BG505 complex, while the W427_gp120_ side chain adopted a position similar to that in the 1-18– BG505 complex. This suggests that, while the overall conformation of the Env trimer in the CD4-HT1 complex represented a closed prefusion Env, the gp120 Phe43 cavity showed indications of structural changes consistent with CD4 binding.

### BG505-HT2 bound to two CD4s adopts an asymmetric, open Env conformation

To structurally characterize BG505-HT2 complexed with CD4, we collected single particle cryo-EM data and recovered three classes that resembled Env-CD4 complexes (Extended Data Fig. 3). Class I (92,660 particles; 3.9 Å resolution) contained a BG505 heterotrimer with two CD4-bound protomers and one unliganded protomer (Extended Data Fig. 3). Class II (48,577 particles; 3.8 Å resolution), which was similar to the CD4-HT1 structure, contained a BG505 trimer bound to a single CD4 (Extended Data Fig. 3). Class III (28,548 particles; 6.4 Å resolution) was poorly resolved (Extended Data Fig. 3), with one Env protomer showing clear density indicative of CD4 binding, while the adjacent protomer showed less defined CD4 density, and density for the third, unliganded protomer extended from the Env gp120 across the trimer apex, appearing to contact the adjacent CD4-bound protomer. Further interpretation was prevented by the limited resolution. Subsequent analyses of the BG505 CD4-HT2 complex were confined to the 3.9 Å class I structure.

We compared the two CD4-bound HT2 structure to other Env conformations by quantifying gp120 rearrangements using measurements of inter-protomer distances between the Cα atoms of conformationally-characteristic Env residues (Fig. 2a,b). The relationship between the HT2 gp120 protomers that bound CD4 resembled a typical CD4-induced open conformation, with V1V2 loops displaced from the Env apex to the sides of gp120 and the V3 loops exposed (Fig. 2a), consistent with increased inter-protomer distances between these protomers compared to closed^33^, occluded-open^12^, and partially open^8^ Env conformations (Fig. 2b). The unliganded HT2 protomer did not show V1V2 and V3 loop movement to the extent observed in the CD4-bound protomers. Instead, the V1V2 and V3 loops were displaced as a rigid body from the Env apex, as observed in the protomers of the homotrimeric occluded-open Env conformation (Fig. 2a)^12^. Asymmetry of the HT2 Env with two bound CD4s was demonstrated by variable inter-protomer distances: the measured distance between the two CD4-bound gp120s (protomers A and B) were consistent with the open, CD4-bound Env conformation, by contrast to distances between the CD4-bound gp120s and the unliganded gp120 (protomer C), which were slightly smaller than distances between CD4-bound gp120s. Thus, the HT2 Env adopted an asymmetric conformation in which the distance to the central trimer axis was smaller in the unliganded protomer than in the CD4-bound protomers (Fig. 2b; Supplementary Movie 1).

**Fig. 2:**
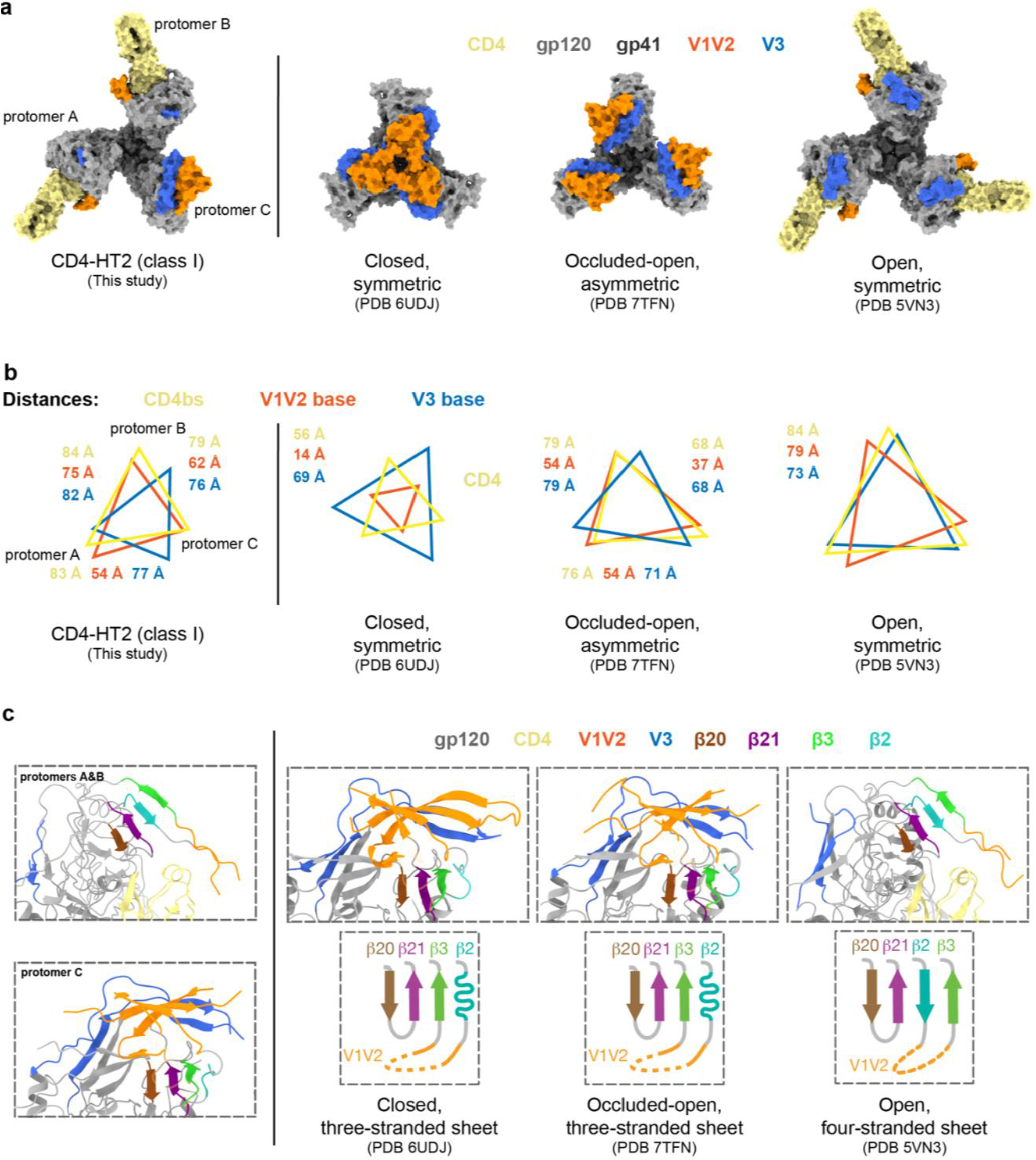
3.9 Å cryo-EM structure of BG505-HT2 bound to two CD4 molecules reveals an asymmetric open Env conformation. **a**, Top-down views of surface depictions of CD4-HT2 (class I), an Env in a prefusion conformation (PDB 6UDJ), Env in the occluded-open conformation (PDB 7TFN), and the CD4-bound open conformation of Env (PDB 5VN3). **b**, Inter-protomer distance measurements between reference residues for the base of the V3 loop (H330_gp120_), the base of the V1/V2 loop (P124_gp120_), and the CD4bs (D/R368gp120) for CD4-HT2 (class I), an Env in a prefusion conformation (PDB 6UDJ), Env in the occluded-open conformation (PDB 7TFN), and the CD4-bound open conformation of Env (PDB 5VN3). **c**, Cartoon representations of the gp120 bridging sheet motif for CD4-HT2 (class I), an Env in a prefusion conformation (PDB 6UDJ), Env in an occluded-open conformation (PDB 7TFN), and the CD4-bound open conformation of Env (PDB 5VN3).

Since a hallmark of CD4-induced gp120 structural changes is the transition of the three-stranded β-sheet to a four-stranded antiparallel bridging sheet^6–8,10^, we next examined the β-sheet conformations in the CD4-HT2 complex. The β-sheet conformations observed in the CD4-HT2 complex differed: CD4-bound protomers A and B included the four-stranded bridging sheet observed in CD4-bound open Env trimer structures^6–8^, whereas the unliganded gp120 in protomer C contained a three-stranded sheet resembling its counterpart gp120s in closed and occluded open conformations^12,33^ (Fig. 2c). In summary, the binding of two CD4s to BG505-HT2 resulted in an asymmetric and partially open Env trimer composed of two CD4-bound, open conformation gp120s and one unliganded gp120 in an occluded-open conformation.

To address the generality of the effects of Env interactions with sub-stoichiometric numbers of CD4s, we prepared HT2 heterotrimers for the clade B B41 SOSIP.664^36^ (Extended Data Fig. 1a), obtaining a 4.1 Å cryo-EM density map of B41-HT2 bound to two CD4 molecules (Extended Data Fig. 4). Fitting the CD4-BG505 HT2 structure into the density map for CD4-B41 HT2 showed agreement in the overall structural features, including V1V2 displacement of CD4-bound protomers and partial outward gp120 rotation of the unliganded protomer (Extended Data Fig. 4). In addition, we solved 4.2 Å and 3.8 Å single particle cryo-EM structures of CD4 complexes with BG505 HT1 and HT2 plus 17b^37^, a CD4-induced antibody that recognizes the exposed coreceptor binding site on V3^6–9^ (Extended Data Fig. 5). For both complexes, the Envs showed three bound 17b Fabs and three bound CD4 molecules and adopted open conformations, as indicated by density for V1V2 that was displaced to the sides of gp120 on each protomer (Extended Data Fig. 5). However, the poor local map density surrounding the Fab-gp120 interface and CD4 prevented building of reliable atomic models.

### gp41 conformational changes are mediated by gp120 conformations in CD4-bound heterotrimer Env structures

HIV-1 gp41 subunits are responsible for fusion events between host and viral membranes to enable infection^1,38,39^. Prefusion gp41 is composed of a long HR1 helix that extends from beneath the gp120 apex, an HR2 helix that surrounds the N-termini of the HR1 coils, and the fusion peptide (FP) and fusion peptide proximal region (FPPR) located between the HR1 and HR2 helices^1,38,40^. CD4 binding leads to compacting of the C-termini of the HR1 (HR1_c_) helices, triggering formation of a pre-hairpin intermediate in which HR1 extends away from HR2 and the viral membrane^1,38,40^. These movements lead to the formation of compact FPPR helices and the transition of the fusion peptides from α-helices that are shielded in hydrophobic environments to solvent-exposed disordered loops^1,10,38,40^.

Previous studies suggested that changes in Env gp120 conformation are correlated with changes in gp41, suggesting cooperativity between the gp120 and gp41 subunits^6–8,10^. Indeed, in closed and CD4-saturated open Env conformations, gp41 subunits undergo the characterized CD4-induced changes described above (Fig. 3a). Closed Env trimers contain gp41s with a disordered HR1_c_, a helical FP, and an FPPR bent helix, while the gp41 subunits in open Env conformations contain a helical HR1_c_, disordered FP, and straight helical FPPR (Fig. 3b). This pattern is also evident in the CD4-HT1 complex, in which despite engagement of one CD4, each of the three gp120 and gp41 subunits mostly retain closed, prefusion conformations (Fig. 3a). The only deviation from the closed gp41 conformation in the HT1 heterotrimer is a disordered FP in all protomers (Fig. 3b).

**Fig. 3:**
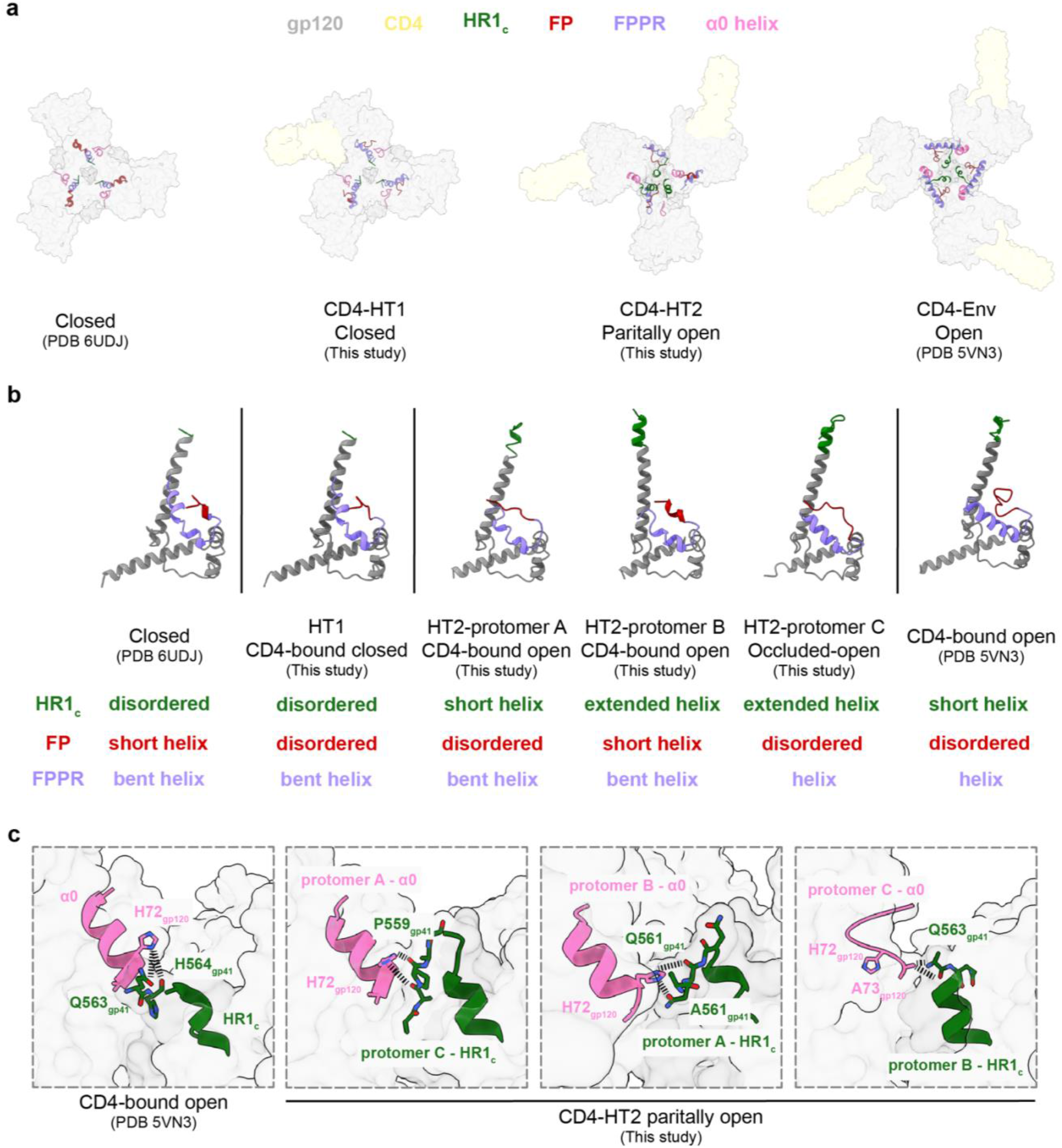
Conformational changes in gp41 were coordinated with gp120 conformation in CD4-bound heterotrimers. **a**, Top-down views of surface representations of Envs from closed (PDB 6UDJ), CD4-HT1, CD4-HT2, and CD4-Env (PDB 5VN3) structures with gp41 structural elements (HR1_c_, FP, FPPR, α0 helix) depicted in cartoon representations. **b**, Cartoon representations of gp41 subunits from closed (PDB 6UDJ), CD4-HT1, CD4-HT2 (protomers a-c), and CD4-Env (PDB 5VN3) structures with colored gp41 (HR1_c_, FP, FPPR, α0 helix) structural elements. **c**, Cartoon representations of the α0 helix and HR1_c_ for CD4-Env (PDB 5VN3) and CD4-HT2 (protomers a-c) with stick representations of selected amino acids. Black dashed connecting lines indicate gp120-gp41 interactions within 6.0 Å.

In the CD4-HT2 complex, individual gp41 subunits adopted distinct conformations despite the near identical conformations of the two CD4-bound gp120s (Fig. 3a,b). The gp41 in CD4-bound protomer A revealed a slanted HR1 helix, a short helical HR1_C_, a disordered FP, and a bent helical FPPR (Fig. 3b). The other CD4-bound gp120 in protomer B contained contrasting elements in gp41: the HR1 and HR1_C_ helices were erect (HR1) or fully extended (HR1_C_), consistent with CD4-induced structural changes (Fig. 3b). By contrast, the FP and FPPR resembled their conformations in closed Envs (Fig. 3b). Despite protomer C being unliganded, its gp41 most resembled the CD4-induced gp41 conformation, with a helical HR1_c_, disordered FP, and a helical FPPR (Fig. 3b). Together, these results demonstrate that individual gp41 subunits can adopt different, distinct conformations in the context of a two CD4-bound Env.

A potential link between gp120 and gp41 Env conformations involves the gp120 α0 region. During Env trimer opening, the HR1_c_ extension displaces the α0 disordered loop located above HR1_c_ in the prefusion conformation and forms a stable α-helix that caps the neighboring gp41 HR1 helix (Fig. 3c)^6,9,10^. In the CD4-HT1 complex, the α0 loops resembled those in the prefusion conformation, whereas the α0 conformations in the CD4-HT2 complex were variable (Fig. 3a,c). Despite only a partial extension of HR1_c_ in CD4-bound protomer A of the HT2 heterotrimer, the gp120 α0 helix was formed and displaced towards the protomer C HR1_c_, where it was stabilized through interactions with the short disordered protomer C HR1_c_ tip (Fig. 3c). Similarly, for CD4-bound protomer B, HR1_c_ extension occurred to form a gp120 α0 helix that interacted with its neighboring protomer A HR1_c_ (Fig. 3c). In unliganded protomer C, the gp120 α0 region remained in the prefusion disordered loop conformation despite extension of its HR1_c_ (Fig. 3c). The loop conformation was likely accommodated because protomer C gp120 does not undergo the full outwards displacement from the Env trimer axis. However, partial outwards rotation of protomer C’s gp120 still enabled interactions with the neighboring protomer B HR1_c_ (Fig. 3c). These inter-protomer interactions between gp120s and gp41s in CD4-HT2 rationalize why each gp41 subunit adopted a distinct conformation, suggesting that formation of the α0 helix is dependent of CD4 occupancy and likely drives gp41 conformational changes.

### Two CD4-bound soluble and membrane-bound Env trimers exhibit similar conformations

Cryo-electron tomography (cryo-ET) and sub-tomogram averaging was used to determine the conformations of membrane-bound Envs complexed with sub-stoichiometric numbers of membrane-bound CD4s^41^. We can therefore compare our higher resolution soluble CD4–soluble heterotrimer Env structures with structures of CD4-Env complexes investigated under more physiological conditions.

Rigid body fitting of the CD4-HT1 model into the cryo-ET/sub-tomogram averaged density of a one CD4-bound Env trimer showed major differences (Extended Data Fig. 6). Unlike the closed Env conformation observed for the soluble CD4-HT1 complex (Fig. 1a), the membrane-bound Env adopted a partially open conformation in response to engagement with a single CD4 in which the CD4-bound protomer appeared to undergo CD4-induced conformational changes consistent with V1V2 displacement (Extended Data Fig. 6)^41^.

By contrast, the two CD4-bound membrane-embedded and soluble Envs adopted similar conformations. Rigid body fitting of the soluble CD4-HT2 structure into the corresponding cryo-ET/sub-tomogram averaged CD4-Env density showed the alignment of the positions of the bound CD4s and Env gp120s (Fig. 4a,b). The displaced V1V2 loops in the CD4-HT2 CD4-bound protomers A and B were clearly matched with density from membrane-embedded Env (Fig. 4c,d), and the partial outward gp120 rotation described in unliganded protomer C in the soluble CD4-Env structure (Fig. 2a,b) also aligned with density for the unliganded protomer in the membrane-bound Env (Fig. 4e). However, the V1V2 and V3 densities were not resolved in the cryo-ET map likely due to flexibility of this region, limiting our comparisons of the V1V2 and V3 regions of the unliganded protomer in membrane-bound Env and soluble Env (Fig. 4e).

**Fig. 4:**
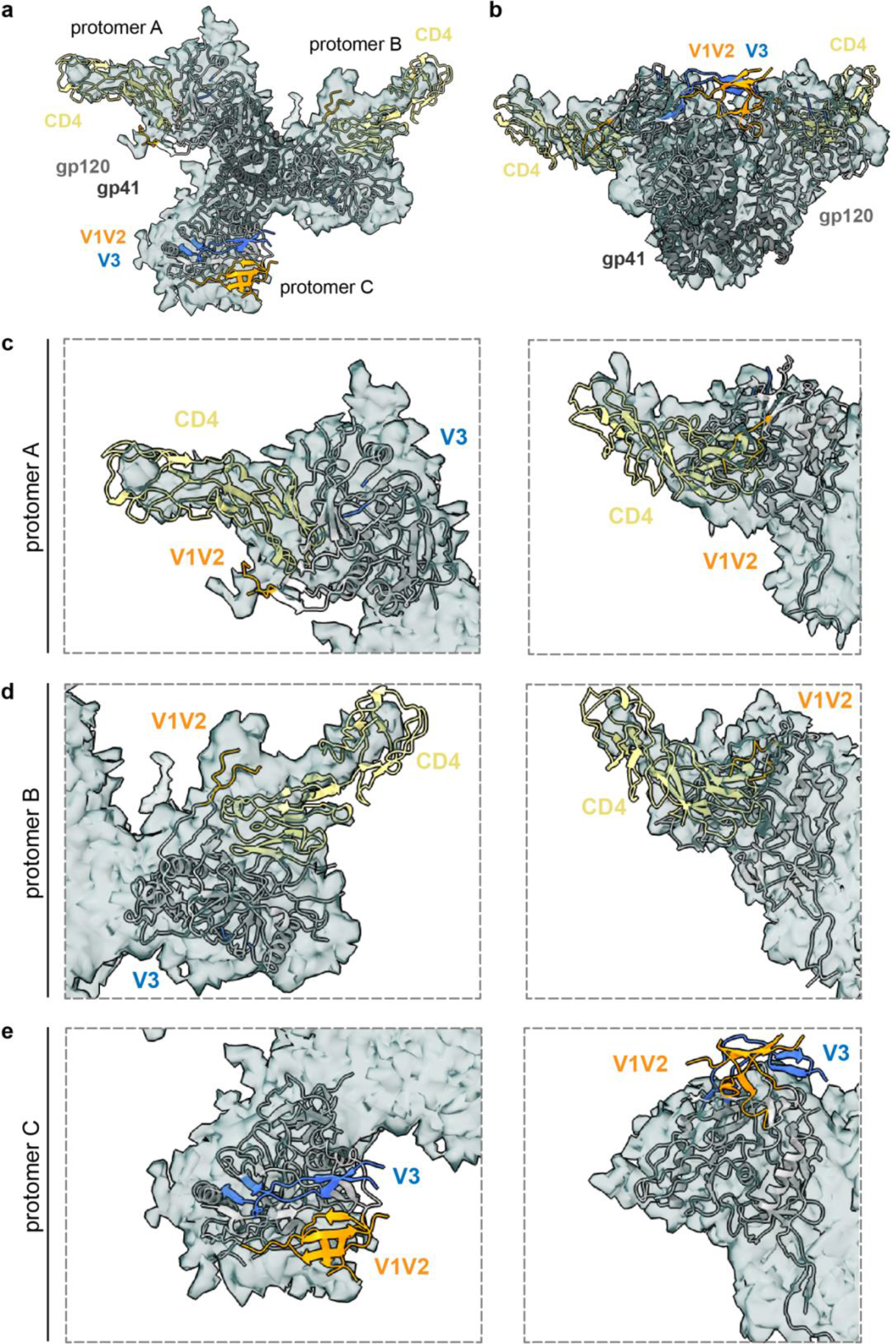
The CD4-HT2 heterotrimer resembles a two CD4-bound membrane-bound Env. **a**, Top-down and **b**, side views of a cartoon representation of the CD4-HT2 (class I) structure fit into density (gray mesh) of a two CD4-bound Env derived from cryo-ET/sub-tomogram averaging^41^. Close-ups of **c**, protomer A, **d**, protomer B, and **e**, protomer C from top-down and side views depicted in **a**, and **b**, respectively.

## Discussion

HIV-1 Env trimers on virions are likely to encounter multiple CD4 receptors on the surface of target cells. However, experimental studies have yet to definitively address whether one, two, or all three CD4 binding sites on each trimer must be occupied for coreceptor binding and downstream conformational changes resulting in viral and host cell membrane fusion. In addition, the degree of cooperativity between Env protomers upon binding to CD4 had not been investigated structurally. The characterization of a non-neutralizing antibody isolated from an immunized macaque that mimicked FP interactions with a single gp41 per trimer, thereby rendering one FP per trimer inactive^42^, implies that not all protomers in each Env trimer are required for virus-host cell membrane fusion. Consistent with this conclusion, fusion and infectivity studies that incorporated Env mutations resulting in defective CD4, coreceptor, and fusion activity in individual protomers of Env heterotrimers^43–46^ suggested that Env entry does not require that each subunit in an individual trimer be competent in performing all functions^43–45^. However, the effects of sub-stoichiometric binding of CD4 in these experiments were complicated by the necessity that Env protomers with different defective mutations were randomly assembled as homotrimeric and heterotrimer Envs that were compared for fusion and infectivity with homotrimeric controls^43–45^. In addition, neither of these types of experiments included structural characterizations to examine the conformational effects of sub-stoichiometric CD4 interactions with individual Env trimers. Thus, our single-particle cryo-EM investigation of Env heterotrimers binding one or two CD4s, together with the accompanying cryo-ET visualization of the native HIV-1 virions and membrane-bound CD4^41^, adds to our knowledge of Env structures, which was previously limited to closed, prefusion Env conformations with either no bound CD4s or three CD4s bound to fully-saturated open Env trimers^2–9,41^.

By engineering soluble Env heterotrimers with either one or two wildtype CD4 binding sites, we solved single-particle cryo-EM structures of Env trimers with sub-stoichiometric numbers of bound CD4s at sufficient resolutions to monitor CD4-induced changes to gp120 and gp41 subunits. We found that binding of one CD4 resulted in minor structural changes to a native-like soluble Env trimer in the closed, prefusion state; for example, we did not observe opening of any of the gp120 subunits of the trimer or the accompanying changes in the CD4-bound gp120 that result from CD4 associating with gp120 in CD4-bound open trimers^6–10^ (in particular, changes resulting from insertion of Phe43_CD4_ into a gp120 hydrophobic cavity, which facilitates induced changes such as V1V2 displacement in CD4-bound gp120 subunits of fully-saturated open Env trimers^6–8,23,24^, were minor). By contrast, the one CD4-bound conformation of membrane-bound Env trimer revealed by cryo-ET/sub-tomogram averaging showed a partially open conformation in which the CD4-bound protomer appeared to undergo CD4-induced conformational changes^41^.

The single-particle cryo-EM and cryo-ET structures of one CD4-bound Env trimers may represent different conformational intermediates involved with engagement of a single CD4, with the closed trimer conformation likely preceding the more open conformation (Fig. 5). Several factors could contribute to the observation of the different one CD4-bound Env trimer conformations: (*i*) Differences in the Env clade being investigated (tier 2 BG505 for single-particle cryo-EM versus tier 1B BaL for cryo-ET), with tier 2 viruses being more resistant to neutralization and likely also CD4-induced changes than tier 1^47^. (*ii*) The increased ability of membrane-bound CD4 compared with soluble CD4 to engage and then dissociate from Envs over the course of an incubation, thus perhaps leading to visualization in the cryo-ET experiments of one CD4-bound Envs that had recently bound two CD4s. (*iii*) SOSIP substitutions that stabilize the prefusion, closed conformation (including the interprotomer disulfide, I556P, A316W)^11,18^ could prevent CD4-induced structural changes when only one CD4 is bound. (*iv*) The CD4-HT1 complex solved by single-particle cryo-EM was prepared at 4°C, whereas the analogous cryo-ET sample was incubated at room temperature – the lower temperature incubation perhaps contributing to observation of the closed trimer conformational state with one CD4 bound that likely precedes a more open trimer conformation (Fig. 5).

**Fig. 5:**
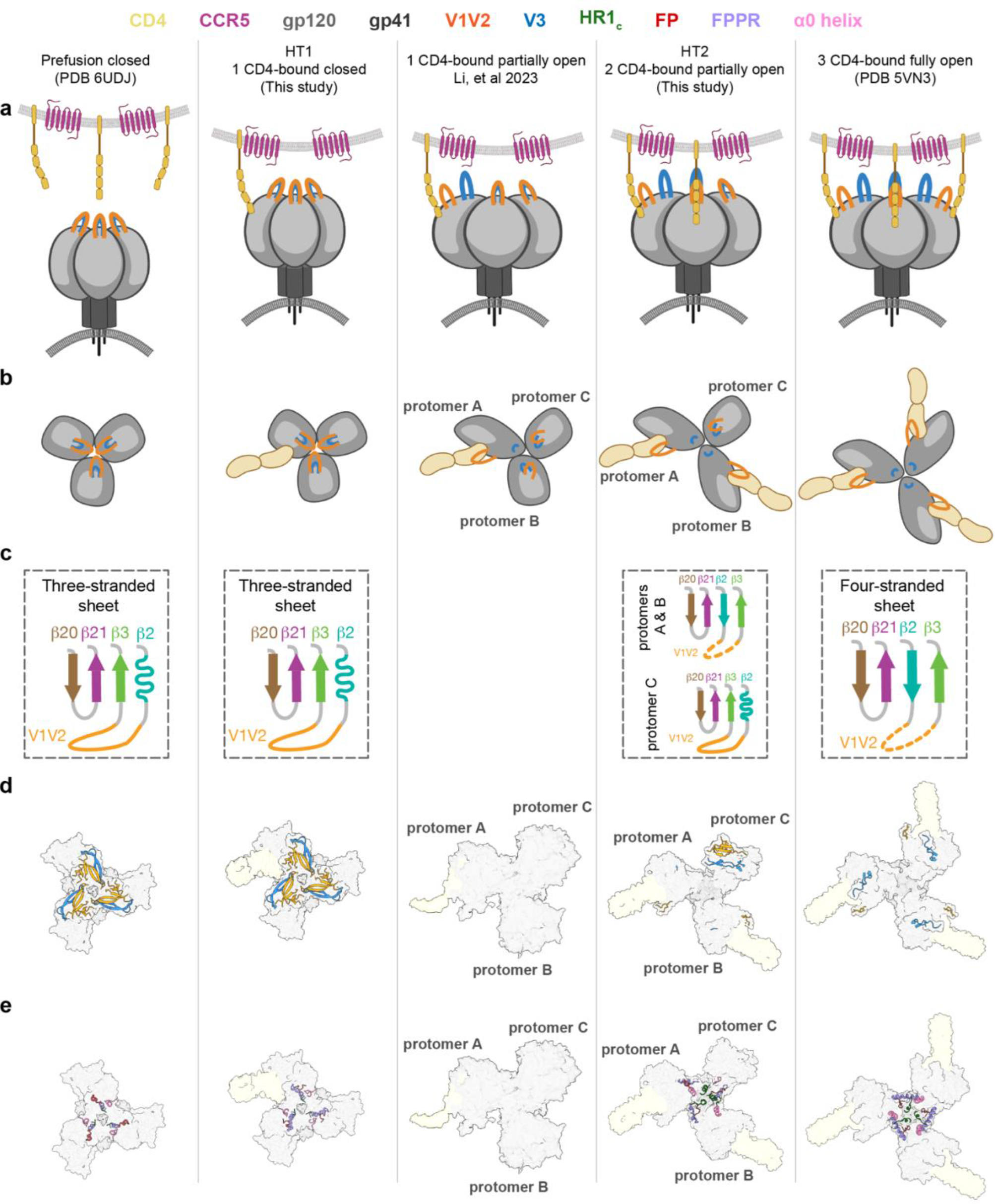
Overview of Env receptor-induced conformational changes that lead to coreceptor binding and fusion. Summary of Env conformational changes, including Envs in prefusion, closed (PDB 6UDJ), one-CD4 bound closed, one-CD4 bound open, two-CD4 bound, partially open, and three-CD4 bound (PDB 5VN3) conformations. Schematics were generated using BioRender. Depictions for each Env conformation include: Env schematics in **a**, side and **b**, top-down views, **c**, diagrams describing β-sheet conformations observed in Env gp120s, **d**, surface representations of structures for each Env conformation with cartoon representations of V1V2 and V3 loops and **e**, gp41 structural features.

The single-particle cryo-EM and cryo-ET structures of two CD4-bound Env trimers were remarkably consistent, such that both showed two protomers in CD4-bound open conformations and the remaining unbound protomer in a conformation resembling an occluded open Env protomer (Fig. 4)^12^. These results provide further evidence of native-like soluble SOSIP Env trimers resembling their virion-bound counterparts^4,5^, both in the closed, prefusion conformation and in various CD4-bound conformations that adopt different conformations compared with unliganded Env trimers. Thus, this and the accompanying cryo-ET study^41^, together with previous Env structures, complete a description of the conformations of HIV-1 Env trimers at each stage of engaging CD4, starting with no bound receptors to the final conformation with three bound receptors (Fig, 5; Supplementary Movie 1).

The ability to confirm single-particle soluble Env heterotrimer conformations that include residue-level details using lower-resolution Env trimer structures derived by cryo-ET under more physiological conditions^41^ lends confidence to the proposed order of structural transitions induced by CD4 binding (Fig. 5; Supplementary Movie 1). The single-particle cryo-EM structures also include the first descriptions of details of CD4-induced structural changes in gp120 and gp41, including cooperative inter-subunit structural transitions. These results reveal intermediate Env conformations that expand our understanding of receptor-induced structural changes preceding host and viral membrane fusion, thereby informing the design of therapeutics to block HIV-1 infection.

## Methods

### Protein expression and purification

SOSIP.664v4 Env constructs included the following stabilizing mutations: introduced cysteines 501C and 605C (SOS), I559P (IP), A316W, and the furin cleavage site mutated to six arginine residues (6R)^11,18^. SOSIPs with D7324 tags included a GSAPTKAKRRVVQREKR sequence after residue 664 in the gp41 ectodomain^11^. The D368R mutation was encoded in Envs to impair CD4 binding^19,48–50^. Genes encoding tagged and untagged SOSIP.664 Env homotrimers were expressed by transient transfection of Expi293 cells (ThermoFisher). Env heterotrimers were purified from co-transfections involving a 20:1 expression plasmid DNA ratio of untagged to tagged Env constructs: a 20:1 ratio of Env-D368R:Env-D7324 (HT1) and a 20:1 plasmid of Env:Env-D368R-D7324 (HT2). Trimeric Envs were purified from cell supernatants by PGT145 immunoaffinity chromatography and size-exclusion chromatography (SEC) using a Superose 6 10/300 column (Cytiva)^11,51^. Tagged Env homotrimers and heterotrimers (HT1 and HT2) were further purified using JR-52 immunoaffinity chromatography as described^11^.

Genes encoding CD4 D1D2 (domains 1 and 2) and D1-D4 (domains 1-4) with C-terminal 6x-His or StrepII tags were transiently transfected using the Expi293 expression system (ThermoFisher)^7^. CD4 proteins were purified using Ni^2+^-NTA (Cytiva) or Strep-Tactin XT (IBA Life Sciences) affinity columns, followed by SEC using a Superdex 200 10/300 column (Cytiva).

The Fab from the CD4i antibody 17b^37^ was expressed by transient transfection using expression vectors encoding the LC and a C-terminally tagged HC portion of the Fab using the Expi293 expression system (ThermoFisher)^7^. Fab was purified from cell supernatants by Ni^2+^-NTA (Cytiva) chromatography followed by SEC using a Superdex 200 10/300 column (Cytiva).

### D7324 capture ELISA

ELISAs were performed as described^9,12,52^. Briefly, 5 μg/mL of JR-52 IgG^11^ (kind gift of James Robinson, Tulane University) was coated on Corning Costar high-binding 96-well plates in 0.1 M NaHCO_3_ (pH 9.6). Plates were incubated overnight at 4 °C. After washing, plates were blocked with 3% BSA in TBS-T (20 mM Tris, 150 mM NaCl, 0.1% Tween-20) for 1 hour at room temperature. Blocking buffer was removed and D7324-tagged Envs were applied to plates at 5 μg/mL in 3% BSA in TBS-T. Plates were incubated for 1 hour at room temperature, and then buffer was removed. For some experiments, 6x-His tagged CD4 was serially diluted in 3% BSA in TBS-T at a top concentration of 100 μg/mL, added to plates, and incubated for 4 hours at room temperature. The CD4 solution was removed and plates were washed with TBS-T, twice. A horseradish peroxidase (HRP) labeled secondary against the His tag (Genscript) was added at a 1:5,000 dilution in 3% BSA in TBS-T. Plates were incubated for 30 minutes, and then washed with TBS-T three times. Colorimetric detection of CD4 binding was accomplished using Ultra TMB-ELISA Substrate Solution (ThermoFisher Scientific) and quenching with 1.0 N HCl. Absorption was measured at 450 nm. Two independent biological replicates (n = 2) were performed for all assays.

### Assembly of protein complexes and cryo-EM sample preparation

The D1-D4 version of CD4 was chosen instead of CD4 D1D2 for structural studies with BG505 HT1 and HT2 to increase particle size. HT1-CD4 and HT2-CD4 complexes were prepared by incubating purified Env heterotrimers with a 1.1x molar excess of CD4 D1-D4 overnight at 4 °C. We attempted CD4-Env incubations at different temperatures (namely 37°C and room temperature) and found that overnight incubation at 4 °C produced the most favorable particle quality when frozen on cryo-EM grids. For HT1-CD4-17b and HT2-CD4-17b complexes, 17b Fab was added prior to grid freezing at a 1.1x molar excess and incubated at 4 °C for 30 minutes. QuantiFoil 300 mesh 1.2/1.3 grids (Electron Microscopy Sciences, Hatfield, PA) were glow discharged with PELCO easiGLOW (Ted Pella, Redding, CA) for 1 min at 20 mA. Fluorinated octylmaltoside solution (Anatrace, Maumee, OH) was added to the protein complex to a final concentration of 0.02% (w/v), and 3 μL of the complex/detergent mixture was applied to glow-discharged grids. A Mark IV Vitrobot (Thermo Fisher Scientific, Waltham, MA) was used to blot grids for 3 seconds with 0 blot force using Whatman No.1 filter paper and 100% humidity at room temperature. Grids were plunge-frozen and vitrified in 100% liquid ethane.

### Cryo-EM sample preparation and data collection

Single particle cryo-EM datasets for HT1-CD4, HT2-CD4, HT1-CD4-17b, and HT2-CD4-17b were collected on a 300 keV Titan Krios (Thermo Fisher Scientific, Waltham, MA) cryo-electron microscope equipped with a K3 direct electron detector camera (Gatan, Pleasanton, CA) using SerialEM^53^ automated data collection software. Movies were recorded with 40 frames at a total dosage of 60 e^−^/Å^2^ using a 3×3 beam image shift pattern with 3 exposures per hole in the super resolution mode, a defocus range of −1 to −3 μm, and pixel size of 0.416 Å.

Data were processed using cryoSPARC^54^. Patch motion correction was applied to each dataset with a binning factor of 2, followed by Patch CTF to estimate CTF parameters. The blob picker with a diameter of 100 to 230 Å was used to pick particles. Particles were extracted and then 2D classified. Particle classes representing the expected complex were selected and used for *ab initio* modeling. The *ab initio* models and corresponding particles that represented the expected complex underwent subsequent rounds of heterogeneous, homogeneous, and non-uniform refinement.

### Structural analyses

Structure figures were created with PyMol (Schrödinger LLC) and UCSF ChimeraX^55^. BSA was calculated using PDBePISA^56^ using a 1.4 Å probe. gp120 BSA was calculated for protein components of gp120 without including glycan coordinates. Due to the low resolution of complexes, interactions were assigned tentatively using the following criteria: hydrogen bonds were assigned as pairwise interactions less than 6.0 Å and with an A-D-H angle >90°, and van der Waals interactions were assigned as distances between atoms that were less than 6.0 Å.

## Acknowledgements

We thank W. Mothes and W. Li, and Z. Qin (Yale University) for sharing cryo-ET data, J. Vielmetter, A. Rorick, K. Storm, and the Protein Expression Center in the Beckman Institute at Caltech for expression assistance, J.E. Robinson (Tulane University) for the JR-52 antibody. Electron microscopy was performed in the Caltech Cryo-EM Center with assistance from S. Chen, and A. DeLaitsch for comments on the manuscript. This work was supported by the NIH P50 AI150464 (P.J.B.), National Institute of Allergy and Infectious Diseases (NIAID) Grant HIVRAD P01 AI100148 (to P.J.B.), and the Bill and Melinda Gates Foundation Collaboration for AIDS Vaccine Discovery (CAVD) grant INV-002143 (P.J.B.). The contents of this publication are solely the responsibility of the authors and do not necessarily represent the official views of NIAID or NIH.

## Author contributions

K.A.D., C.F., and P.J.B. designed the research. K.A.D. designed Env constructs, performed protein purification, and conducted ELISAs. C.F. collected structural data. Z.Y. created Supplementary Movie 1. K.A.D., C.F., Z.Y., and P.J.B. analyzed results. K.A.D. and P.J.B. wrote the manuscript with input from co-authors.

## Data availability

The cryo-EM maps and atomic structures have been deposited in the PDB and/or Electron Microscopy Data Bank (EMDB) under accession codes 8FYI [http://doi.org/10.2210/pdb8fyi/pdb] and EMD-29579 [https://www.ebi.ac.uk/pdbe/entry/emdb/EMD-29579] for CD4-BG505 HT1, 8FYJ [http://doi.org/10.2210/pdb8fyj/pdb] and EMD-29580 [https://www.ebi.ac.uk/pdbe/entry/emdb/EMD-29580] for CD4-BG505 HT2 (class I), EMD-29581 [https://www.ebi.ac.uk/pdbe/entry/emdb/EMD-29581] for CD4-BG505 HT2 (class II), EMD-29582 [https://www.ebi.ac.uk/pdbe/entry/emdb/EMD-29582] for CD4-BG505 HT2 (class III), EMD-29601 [https://www.ebi.ac.uk/pdbe/entry/emdb/EMD-29601] for CD4-B41 HT2, EMD-29583 [https://www.ebi.ac.uk/pdbe/entry/emdb/EMD-29583] for CD4-17b-BG505 HT1, and EMD-29584 [https://www.ebi.ac.uk/pdbe/entry/emdb/EMD-29584] for CD4-17b-BG505 HT2.

## References

1. Harrison, S. C. Viral membrane fusion. Virology 479–480, 498–507 (2015).

2. Lyumkis, D. et al. Cryo-EM Structure of a Fully Glycosylated Soluble Cleaved HIV-1 Envelope Trimer. Science 342, 1484–1490 (2013).

3. Julien, J.-P. et al. Crystal Structure of a Soluble Cleaved HIV-1 Envelope Trimer. Science 342, 1477–1483 (2013).

4. Liu, J., Bartesaghi, A., Borgnia, M. J., Sapiro, G. & Subramaniam, S. Molecular architecture of native HIV-1 gp120 trimers. Nature 455, 109–113 (2008).

5. Li, Z. et al. Subnanometer structures of HIV-1 envelope trimers on aldrithiol-2-inactivated virus particles. Nat. Struct. Mol. Biol. 27, 726–734 (2020).

6. Ozorowski, G. et al. Open and closed structures reveal allostery and pliability in the HIV-1 envelope spike. Nat. Publ. Group 547, 360–363 (2017).

7. Wang, H. et al. Cryo-EM structure of a CD4-bound open HIV-1 envelope trimer reveals structural rearrangements of the gp120 V1V2 loop. Proc. Natl. Acad. Sci. U. S. A. 113, E7151–E7158 (2016).

8. Wang, H., Barnes, C. O., Yang, Z., Nussenzweig, M. C. & Bjorkman, P. J. Partially Open HIV-1 Envelope Structures Exhibit Conformational Changes Relevant for Coreceptor Binding and Fusion. Cell Host Microbe 24, 579–592.e4 (2018).

9. Jette, C. A. et al. Cryo-EM structures of HIV-1 trimer bound to CD4-mimetics BNM-III-170 and M48U1 adopt a CD4-bound open conformation. Nat. Commun. 12, 1950 (2021).

10. Yang, Z., Wang, H., Liu, A. Z., Gristick, H. B. & Bjorkman, P. J. Asymmetric opening of HIV-1 Env bound to CD4 and a coreceptor-mimicking antibody. Nat. Struct. Mol. Biol. 26, 1167–1175 (2019).

11. Sanders, R. W. et al. A Next-Generation Cleaved, Soluble HIV-1 Env Trimer, BG505 SOSIP.664 gp140, Expresses Multiple Epitopes for Broadly Neutralizing but Not Non-Neutralizing Antibodies. PLoS Pathog. 9, e1003618–20 (2013).

12. Yang, Z. et al. Neutralizing antibodies induced in immunized macaques recognize the CD4-binding site on an occluded-open HIV-1 envelope trimer. Nat. Commun. 13, 732 (2022).

13. Stadtmueller, B. M. et al. DEER Spectroscopy Measurements Reveal Multiple Conformations of HIV-1 SOSIP Envelopes that Show Similarities with Envelopes on Native Virions. Immunity 1–27 (2018) doi:10.1016/j.immuni.2018.06.017.

14. Pan, J., Peng, H., Chen, B. & Harrison, S. C. Cryo-EM Structure of Full-length HIV-1 Env Bound With the Fab of Antibody PG16. J. Mol. Biol. 432, 1158–1168 (2020).

15. Harris, A. et al. Trimeric HIV-1 glycoprotein gp140 immunogens and native HIV-1 envelope glycoproteins display the same closed and open quaternary molecular architectures. Proc. Natl. Acad. Sci. 108, 11440–11445 (2011).

16. Gristick, H. B. et al. Natively glycosylated HIV-1 Env structure reveals new mode for antibody recognition of the CD4-binding site. Nat. Publ. Group 23, 906–915 (2016).

17. Scharf, L. et al. Broadly Neutralizing Antibody 8ANC195 Recognizes Closed and Open States of HIV-1 Env. Cell 1–13 (2018) doi:10.1016/j.cell.2015.08.035.

18. de Taeye, S. W. et al. Immunogenicity of Stabilized HIV-1 Envelope Trimers with Reduced Exposure of Non-neutralizing Epitopes. Cell 163, 1702–1715 (2015).

19. Sanders, R. W. et al. HIV-1 neutralizing antibodies induced by native-like envelope trimers. Science 349, aac4223 (2015).

20. Olshevsky, U. et al. Identification of individual human immunodeficiency virus type 1 gp120 amino acids important for CD4 receptor binding. J. Virol. 64, 5701–5707 (1990).

21. Lu, M. et al. Associating HIV-1 envelope glycoprotein structures with states on the virus observed by smFRET. Nature 15, 690–21 (2019).

22. Liu, Q. et al. Quaternary contact in the initial interaction of CD4 with the HIV-1 envelope trimer. Nat. Struct. Mol. Biol. 24, 370–378 (2017).

23. Prévost, J. et al. The HIV-1 Env gp120 Inner Domain Shapes the Phe43 Cavity and the CD4 Binding Site. mBio 11, e00280–20 (2020).

24. Kwong, P. D. et al. Structure of an HIV gp120 envelope glycoprotein in complex with the CD4 receptor and a neutralizing human antibody. Nature 393, 648–659 (1998).

25. Huang, C. et al. Scorpion-Toxin Mimics of CD4 in Complex with Human Immunodeficiency Virus gp120. Structure 13, 755–768 (2005).

26. Stricher, F. et al. Combinatorial Optimization of a CD4-Mimetic Miniprotein and Cocrystal Structures with HIV-1 gp120 Envelope Glycoprotein. J. Mol. Biol. 382, 510–524 (2008).

27. Vita, C. et al. Rational engineering of a miniprotein that reproduces the core of the CD4 site interacting with HIV-1 envelope glycoprotein. Proc. Natl. Acad. Sci. 96, 13091–13096 (1999).

28. Melillo, B. et al. Small-Molecule CD4-Mimics: Structure-Based Optimization of HIV-1 Entry Inhibition. ACS Med. Chem. Lett. 7, 330–334 (2016).

29. Haim, H. et al. Soluble CD4 and CD4-Mimetic Compounds Inhibit HIV-1 Infection by Induction of a Short-Lived Activated State. PLoS Pathog. 5, e1000360 (2009).

30. Courter, J. R. et al. Structure-Based Design, Synthesis and Validation of CD4-Mimetic Small Molecule Inhibitors of HIV-1 Entry: Conversion of a Viral Entry Agonist to an Antagonist. Acc. Chem. Res. 47, 1228–1237 (2014).

31. West Jr, A. P. Structural basis for germ-line gene usage of a potent class of antibodies targeting the CD4-binding site of HIV-1 gp120. PNAS 1–8 (2012) doi:10.1073/pnas.1208984109/-/DCSupplemental.

32. Huang, J. et al. Identification of a CD4-Binding-Site Antibody to HIV that Evolved Near-Pan Neutralization Breadth. Immunity 45, 1108–1121 (2016).

33. Schommers, P. et al. Restriction of HIV-1 Escape by a Highly Broad and Potent Neutralizing Antibody. Cell 180, 471–489.e22 (2020).

34. Barnes, C. O. et al. A naturally arising broad and potent CD4-binding site antibody with low somatic mutation. Sci. Adv. 8, eabp8155 (2022).

35. Zhou, T. et al. Structural Basis for Broad and Potent Neutralization of HIV-1 by Antibody VRC01. Science 329, 811–817 (2010).

36. Pugach, P. et al. A Native-Like SOSIP.664 Trimer Based on an HIV-1 Subtype B *env* Gene. J. Virol. 89, 3380–3395 (2015).

37. Sullivan, N. et al. CD4-Induced Conformational Changes in the Human Immunodeficiency Virus Type 1 gp120 Glycoprotein: Consequences for Virus Entry and Neutralization. J. Virol. 72, 4694–4703 (1998).

38. Chan, D. C. & Kim, P. S. HIV Entry and Its Inhibition. Cell 93, 681–684, (1998).

39. Chan, D. C., Fass, D., Berger, J. M. & Kim, P. S. Core Structure of gp41 from the HIV Envelope Glycoprotein. Cell 89, 263–273 (1997).

40. Ladinsky, M. S. et al. Electron tomography visualization of HIV-1 fusion with target cells using fusion inhibitors to trap the pre-hairpin intermediate. eLife 9, e58411 (2020).

41. Li, W. et al. Asymmetric HIV-1 envelope trimers bound to one and two CD4 molecules are intermediates during membrane binding. http://biorxiv.org/lookup/doi/10.1101/2022.12.23.521843 (2022) doi:10.1101/2022.12.23.521843.

42. Abernathy, M. E. et al. Antibody elicited by HIV-1 immunogen vaccination in macaques displaces Env fusion peptide and destroys a neutralizing epitope. Npj Vaccines 6, 126 (2021).

43. Yang, X., Kurteva, S., Ren, X., Lee, S. & Sodroski, J. Subunit Stoichiometry of Human Immunodeficiency Virus Type 1 Envelope Glycoprotein Trimers during Virus Entry into Host Cells. J. Virol. 80, 4388–4395 (2006).

44. Salzwedel, K. & Berger, E. A. Complementation of diverse HIV-1 Env defects through cooperative subunit interactions: a general property of the functional trimer. Retrovirology 6, 75 (2009).

45. Salzwedel, K. & Berger, E. A. Cooperative subunit interactions within the oligomeric envelope glycoprotein of HIV-1: Functional complementation of specific defects in gp120 and gp41. Proc. Natl. Acad. Sci. 97, 12794–12799 (2000).

46. Khasnis, M. D., Halkidis, K., Bhardwaj, A. & Root, M. J. Receptor Activation of HIV-1 Env Leads to Asymmetric Exposure of the gp41 Trimer. PLOS Pathog. 12, e1006098 (2016).

47. Seaman, M. S. et al. Tiered Categorization of a Diverse Panel of HIV-1 Env Pseudoviruses for Assessment of Neutralizing Antibodies. J. Virol. 84, 1439–1452 (2010).

48. Li, Y. et al. Broad HIV-1 neutralization mediated by CD4-binding site antibodies. Nat. Med. 13, 1032–1034 (2007).

49. Umotoy, J. et al. Rapid and Focused Maturation of a VRC01-Class HIV Broadly Neutralizing Antibody Lineage Involves Both Binding and Accommodation of the N276-Glycan. Immunity 51, 141–154.e6 (2019).

50. Ma, X. & Mothes, W. HIV-1 Env trimer opens through an asymmetric intermediate in which individual protomers adopt distinct conformations. eLife 7:e34271, 1–18 (2018).

51. Cupo, A. et al. Optimizing the production and affinity purification of HIV-1 envelope glycoprotein SOSIP trimers from transiently transfected CHO cells. PLOS ONE 14, e0215106 (2019).

52. Dam, K.-M. A., Mutia, P. S. & Bjorkman, P. J. Comparing methods for immobilizing HIV-1 SOSIPs in ELISAs that evaluate antibody binding. Sci. Rep. 12, 11172 (2022).

53. Mastronarde, D. N. SerialEM: A Program for Automated Tilt Series Acquisition on Tecnai Microscopes Using Prediction of Specimen Position. Microsc. Microanal. 9, 1182–1183 (2003).

54. Punjani, A., Rubinstein, J. L., Fleet, D. J. & Brubaker, M. A. cryoSPARC: algorithms for rapid unsupervised cryo-EM structure determination. Nat. Methods 14, 290–296 (2017).

55. Goddard, T. D. et al. UCSF ChimeraX: Meeting modern challenges in visualization and analysis: UCSF ChimeraX Visualization System. Protein Sci. 27, 14–25 (2018).

56. Krissinel, E. & Henrick, K. Inference of Macromolecular Assemblies from Crystalline State. J. Mol. Biol. 372, 774–797 (2007).

57. Kyte, J. & Doolittle, R. F. A simple method for displaying the hydropathic character of a protein. J. Mol. Biol. 157, 105–132 (1982).

