## Extended Data for "Structural characterization of HIV-1 Env heterotrimers bound to one or two CD4 receptors reveals intermediate Env conformations"

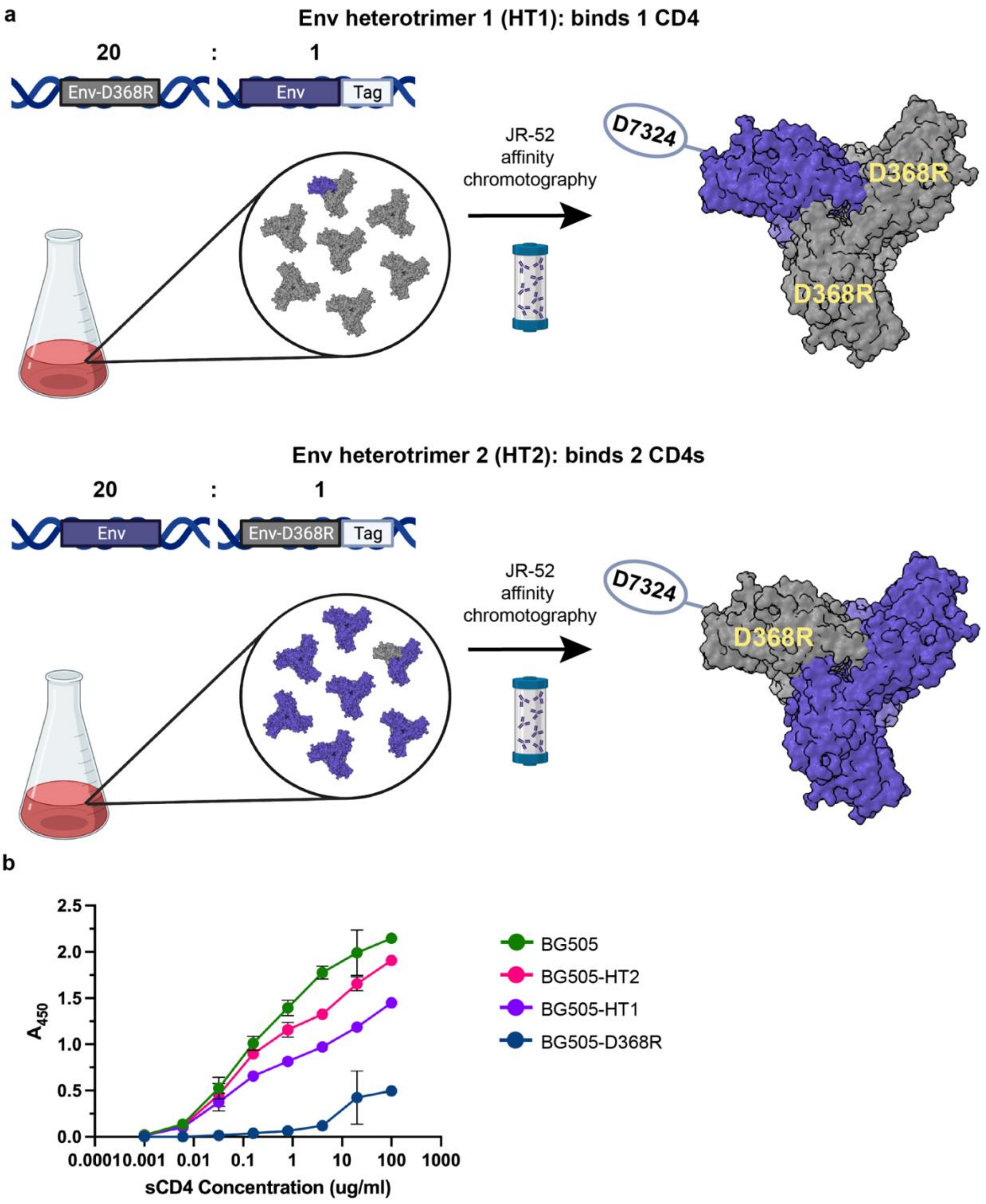


**Extended Data Fig. 1: Design and validation of soluble HIV-1 Env heterotrimer constructs**

**a,** Methods used to create soluble HT1 and HT2 HIV-1 Env heterotrimers. A 20:1 transfection ratio of untagged and D7324-tagged Env expression plasmids, one of which encoded the

D368R CD4 knockout mutation in gp120, was co-transfected to produce two predominant populations: untagged trimers and singly tagged trimers. Transfection supernatants were harvested and Env proteins purified by JR-52 immunoaffinity chromatography (as described), resulting in the HT1 and HT2 heterotrimers. **b,** ELISA comparing CD4 binding of BG505, BG505-HT2, BG505-HT1, and BG505-D368R. Values are shown as mean ± s.d. of two individual biological replicates (n=2). Error bars are not visible for data points where bars are smaller than the size of the symbol representing the mean value.

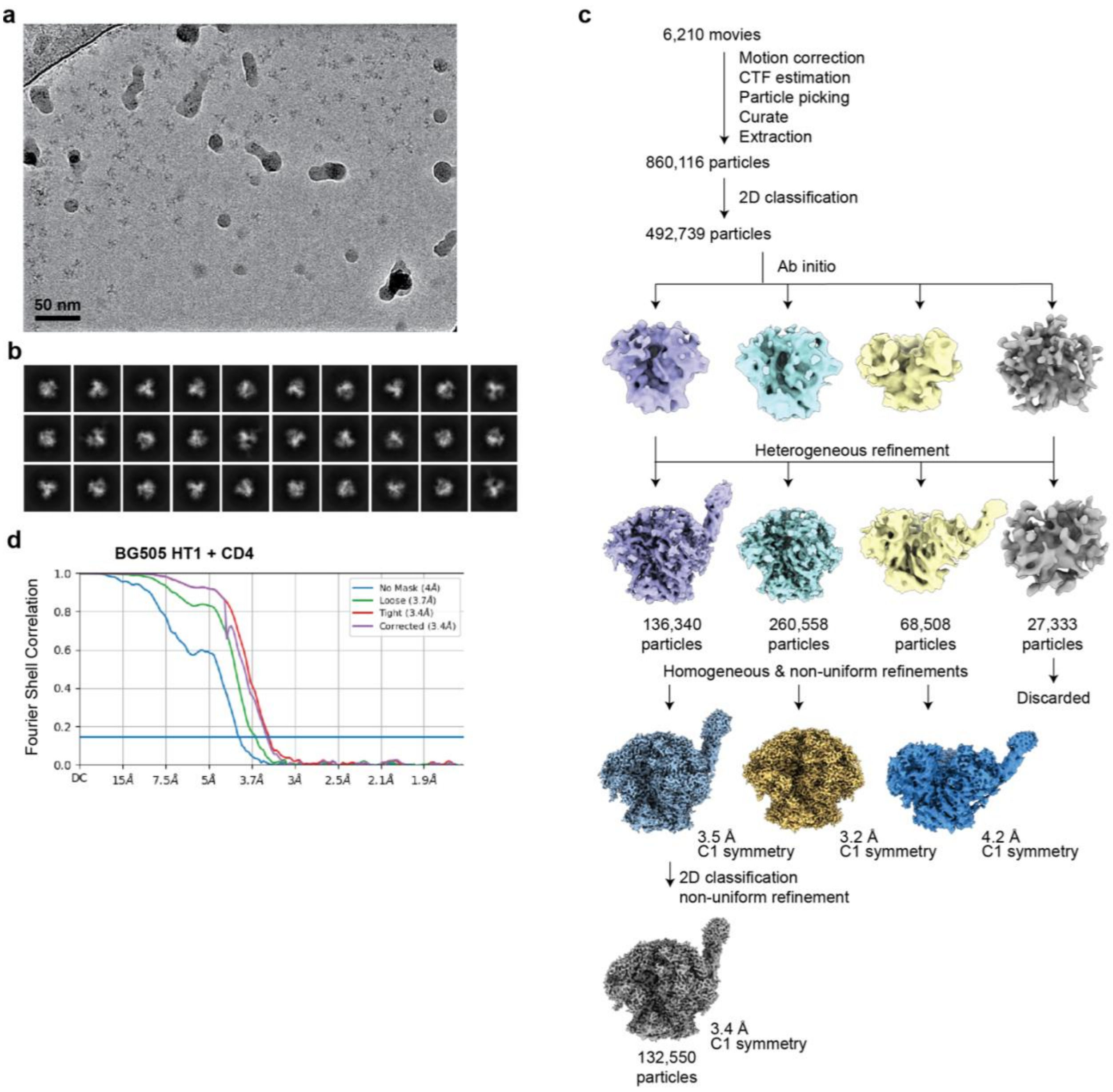


**Extended Data Fig. 2: Cryo-EM data processing and validation for BG505-HT1 in complex with CD4**

**a**, Representative micrograph and **b**, representative 2D classes for the CD4-BG505 HT1 complex. **c**, Workflow of single particle cryo-EM data processing. **d**, Fourier shell correlation (FSC) plot of the final reconstruction for CD4-BG505 HT1.

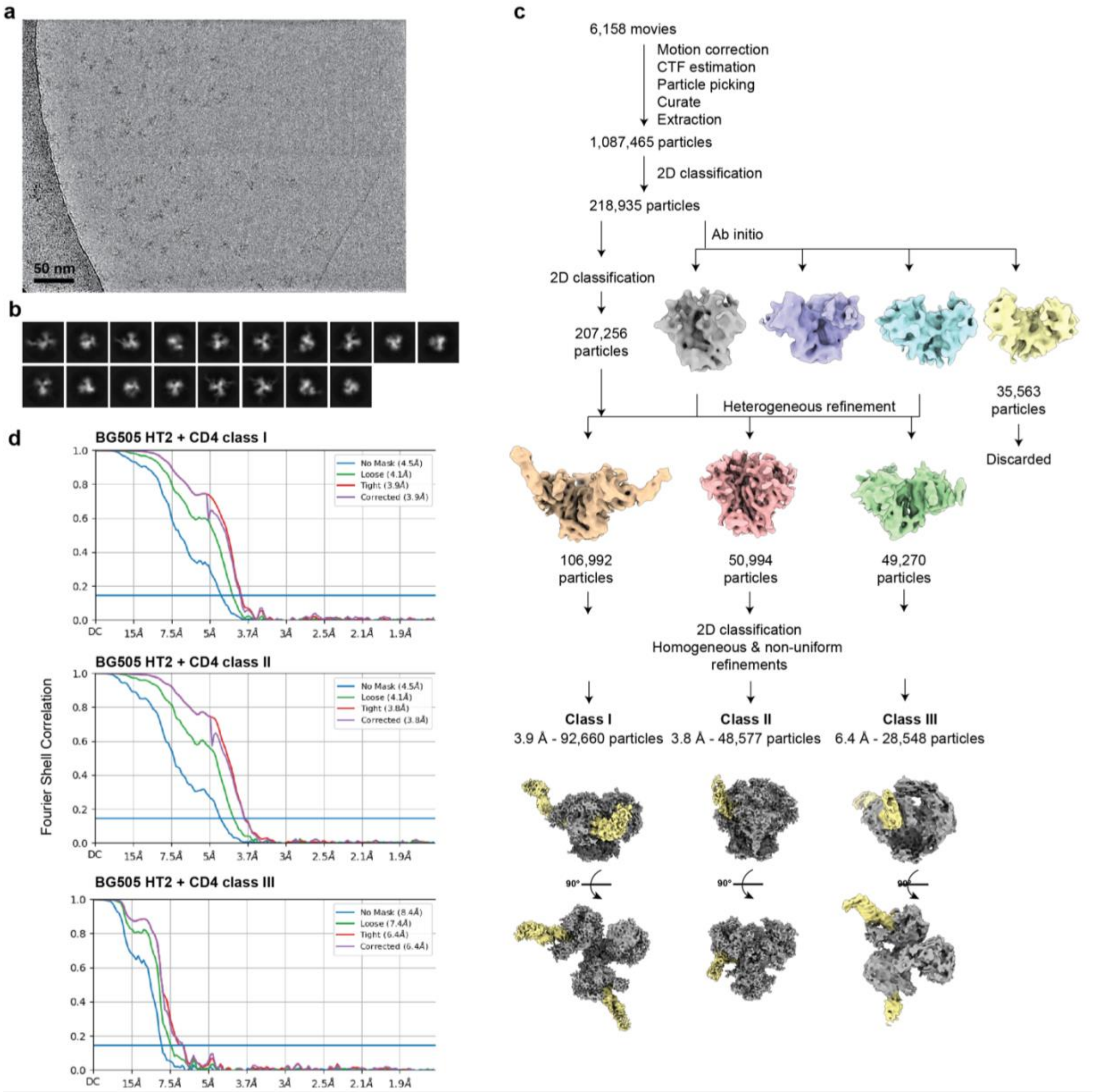


**Extended Data Fig. 3: Cryo-EM data processing and validation for BG505-HT2 in complex with CD4**

**a**, Representative micrograph and **b**, representative 2D classes for the CD4-BG505 HT2 complex. **c**, Workflow of single particle cryo-EM data processing. **d**, Fourier shell correlation (FSC) plot of the final reconstruction for CD4-BG505 HT2 classes I, II, and III.

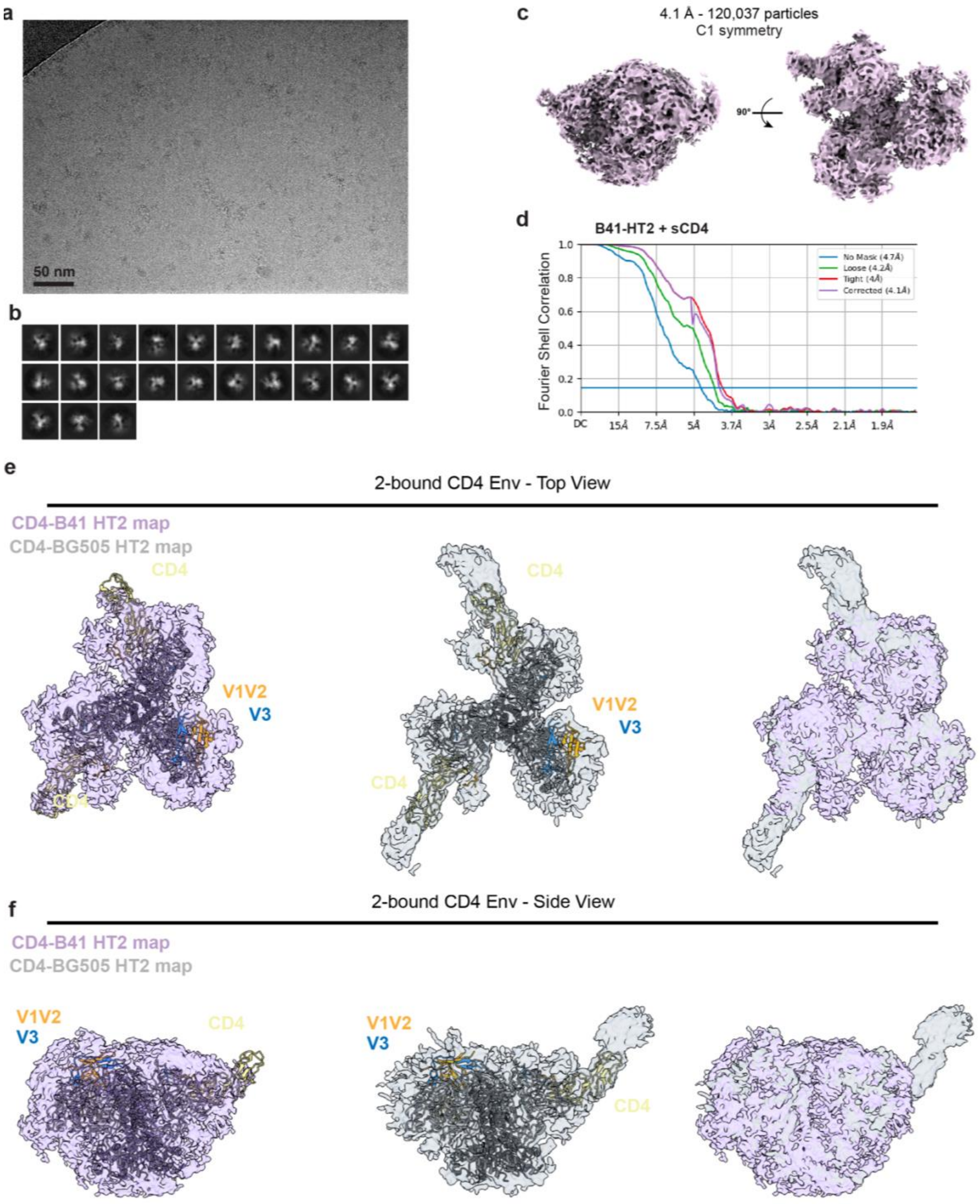


**Extended Data Fig. 4: Cryo-EM data processing, validation, and interpretation for B41-HT2 in complex with CD4**

**a**, Representative micrograph, **b**, representative 2D classes, and **c**, density map for the CD4-B41 HT2 complex. **d**, Fourier shell correlation (FSC) plot of the final reconstruction for CD4-B41 HT2. **e**, Top-down views of CD4-BG505 HT2 model fit into CD4-B41 HT2 (left) or CD4-BG505 HT2 (middle) density maps and alignment of both density maps (right). **f**, Side views of CD4-BG505 HT2 model fit into CD4-B41 HT2 (left) or CD4-BG505 HT2 (middle) density maps and alignment of both density maps (right).

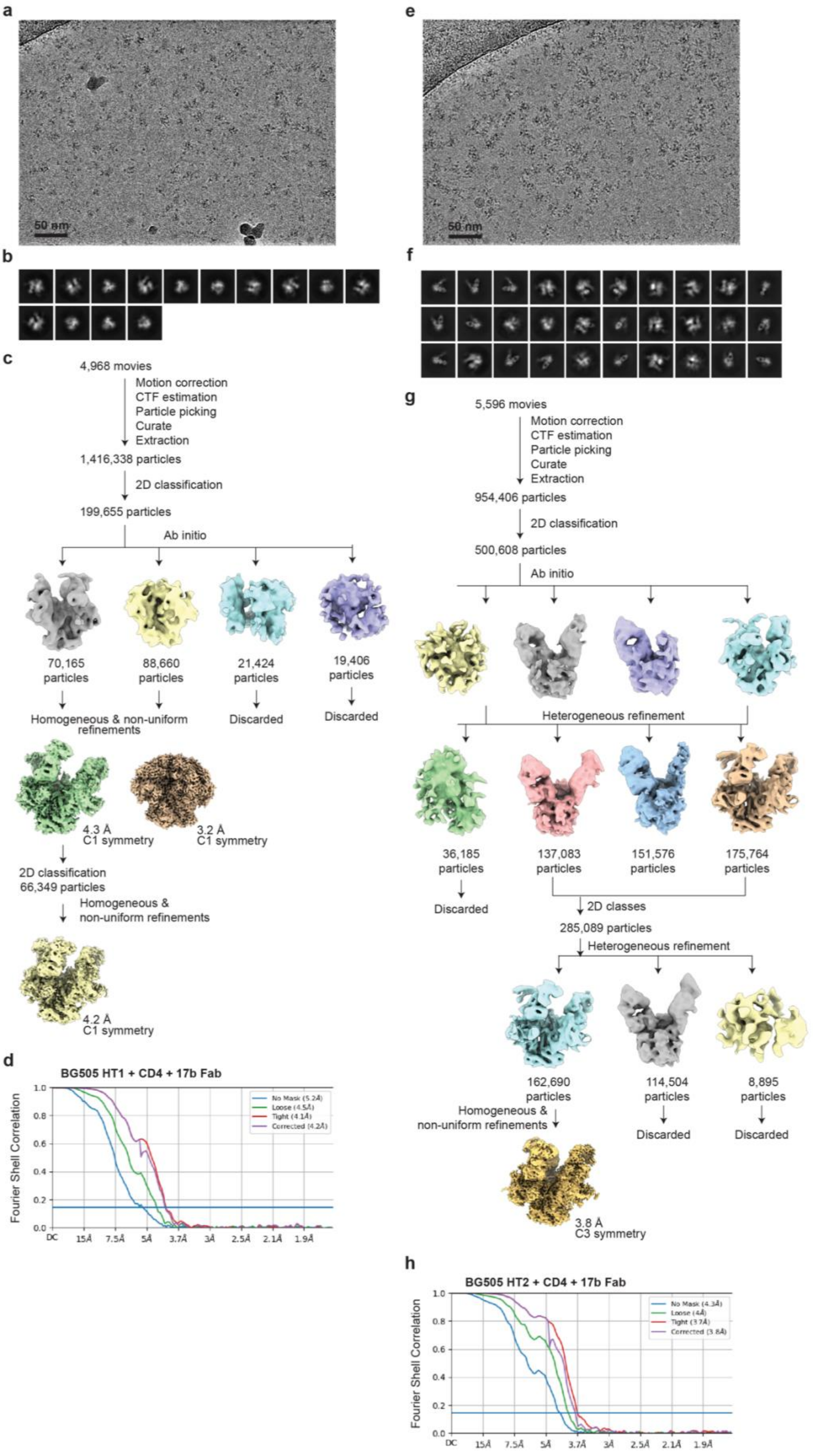

a
50 nm
b
c
4,968 movies
Motion correction
CTF estimation
Particle picking
Curate
Extraction
1,416,338 particles
2D classification
199,655 particles
Ab initio
70,165 particles
88,660 particles
21,424 particles
19,406 particles
Homogeneous & non-uniform refinements
Discarded
Discarded
4.3 Å C1 symmetry
3.2 Å C1 symmetry
2D classification
66,349 particles
Homogeneous & non-uniform refinements
4.2 Å C1 symmetry
d
BG505 HT1 + CD4 + 17b Fab
Fourier Shell Correlation
No Mask (5.2Å)
Loose (4.5Å)
Tight (4.1Å)
Corrected (4.2Å)
DC 15Å 7.5Å 5Å 3.7Å 3Å 2.5Å 2.1Å 1.9Å
e
50 nm
f
g
5,596 movies
Motion correction
CTF estimation
Particle picking
Curate
Extraction
954,406 particles
2D classification
500,608 particles
Ab initio
Heterogeneous refinement
36,185 particles
137,083 particles
151,576 particles
175,764 particles
Discarded
2D classes
285,089 particles
Heterogeneous refinement
162,690 particles
114,504 particles
8,895 particles
Homogeneous & non-uniform refinements
Discarded
Discarded
3.8 Å C3 symmetry
h
BG505 HT2 + CD4 + 17b Fab
Fourier Shell Correlation
No Mask (4.3Å)
Loose (4Å)
Tight (3.7Å)
Corrected (3.8Å)
DC 15Å 7.5Å 5Å 3.7Å 3Å 2.5Å 2.1Å 1.9Å

**Extended Data Fig. 5: Cryo-EM data processing and validation for BG505-HT1 and BG505-HT2 in complex with CD4 and 17b Fab**

**a**, Representative micrograph and **b**, representative 2D classes for the CD4-17b-BG505 HT1 complex. **c**, Workflow of single particle cryo-EM data processing. **d**, Fourier shell correlation (FSC) plot of the final reconstruction for CD4-17b-BG505 HT1. **e**, Representative micrograph and **f**, representative 2D classes for the CD4-17b-BG505 HT2 complex. **g**, Workflow of single particle cryo-EM data processing. **h**, Fourier shell correlation (FSC) plot of the final reconstruction for CD4-17b-BG505 HT2.

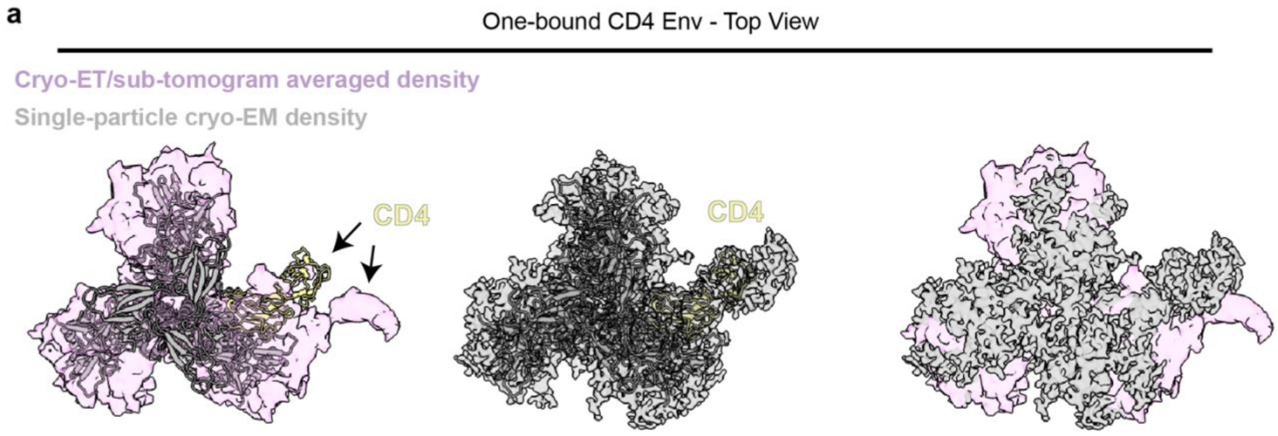


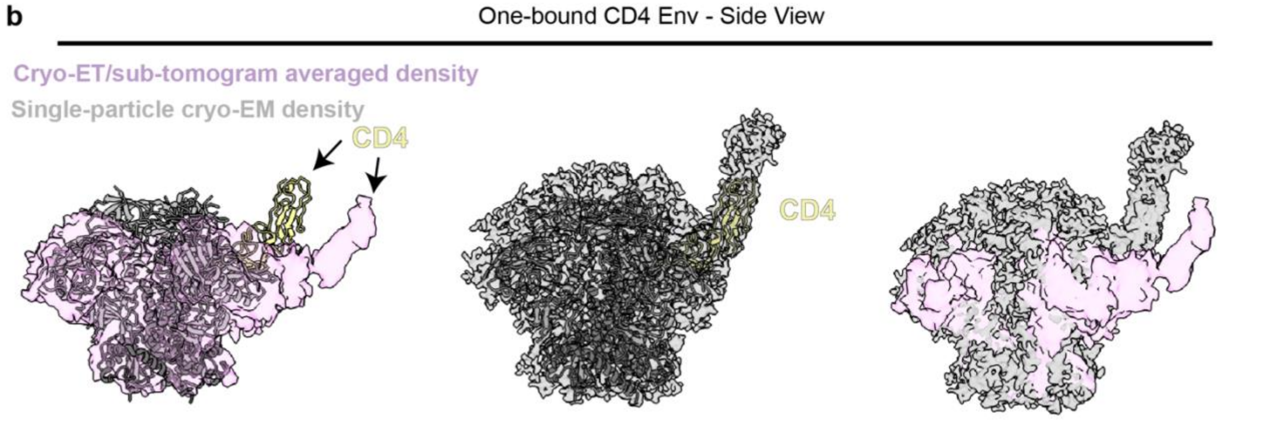


**Extended Data Fig. 6: Comparisons of one CD4-bound Env cryo-ET/sub-tomogram averaged density with CD4-HT1 density and model**

**a**, Top-down views of CD4-HT1 model fit into one-CD4 bound Env from cryo-ET/sub-tomogram averaged (left) or single-particle cryo-EM (middle) density maps and alignment of cryo-ET/sub-tomogram averaged and single-particle cryo-EM density maps (right). **b**, Side views of CD4-HT1 model fit into one-CD4 bound Env from cryo-ET/sub-tomogram averaged (left) or single-particle cryo-EM (middle) density maps and alignment of cryo-ET/sub-tomogram averaged and single-particle cryo-EM density maps (right).
